# Molecular dynamics simulations demonstrate reduced antibiotic affinity to mirror bacterial targets

**DOI:** 10.64898/2025.12.17.694932

**Authors:** Paul-Enguerrand Fady, JL Ciccone

## Abstract

“Mirror life”, self-replicating organisms composed of non-natural-chirality biomacromolecules, presents a severe future threat with potentially global consequences. Consequently, there is strong agreement among experts that it should not be created. However, there is some disagreement over how effective existing medical countermeasures might prove against mirror bacteria in the event that they were created. Here, we leverage computational chemistry methods including docking and molecular dynamics to determine the likely binding efficacy of existing antibiotics against natural and mirror bacterial protein targets. We find that existing most antibiotics fail to bind to mirror bacterial protein targets, unlike their natural chirality targets. This suggests that current medical countermeasures would not successfully exert an antimicrobial activity against mirror bacteria if the latter were created. Our results motivate further policy advocacy to curtail research that directly leads to the creation of mirror life.

## Introduction

Living organisms are composed of biomacromolecules with a specific chirality. All known life forms exhibit homochirality: glycans are predominantly composed of sugars in the D-configuration (such as D-glucose, D-mannose, D-galactose), i.e. sugars with right-handed chiral centres.^1^ The D-2-deoxyribose of DNA is also chiral, imparting a right-handed directionality to the helical twist.^2^ Conversely, proteins in living organisms are composed predominantly of L-amino acids, i.e. ones with left-handed chiral centres.^3^ While it is possible for monomers of the opposite chirality to be integrated into biomacromolecules (and even common in *e*.*g*. peptidoglycan bacterial cell walls which contains D-alanine and D-glutamate),^4^ the predominant pattern of chirality is fixed in one specific form for individual biomacromolecules.

“Mirror life”, or synthetic living organisms composed of biomacromolecules built using the non-naturally occurring enantiomers of biomolecular monomers, presents an alternative option to this universal truth. Recent advances in synthetic cell biology have transformed mirror life from a theoretical science-fiction scenario to a material future threat. While the exact timeline for the development of self-replicating mirror organisms is disputed, experts estimate that it could be achieved within the next 10 to 30 years, depending on resource allocation.^5^

Mirror life could represent a severe future threat with potentially global consequences. A technical report published in December 2024 outlines the ways in which mirror bacteria could subvert human, animal, and plant immune systems, leading to uncontrolled growth due to lack of recognition by stereospecific defences.^5^ Were mirror bacteria to be created and find their way into the environment – whether through deliberate release of a containment breach – they could proliferate extensively. This could ultimately lead to nutrition depletion in key ecological niches, starving keystone species of nutrients and causing ecological collapse. On a wider scale, the sequestration of nutrients from biogeochemical flows would lead to widespread environmental collapse.

The environmental picture motivates policy recommendations to ban research that directly leads to the creation of mirror life. This is distinct from research which may involve mirror macromolecules, which must be permitted in order to foster innovation in the life and material sciences. There is widespread agreement among scientific experts from a range of domains that mirror life should not be created, as noted in the Policy Forum piece accompanying the 2024 technical report;^5^ the “Spirit of Asilomar” conference statement on mirror life in early 2025;^6^ the Paris Conference on Risks from Mirror Life Meeting Report from mid-2025;^7^ a commentary by the German Central Commission for Biological Safety;^8^ and a report from the UNESCO International Bioethics Committee.^9^

However, there has been pushback from some experts surrounding the actual level of risk associated with self-replicating mirror organisms, such as mirror bacteria. A small minority of dissenting voices have indicated their belief that current medical countermeasures such as antibiotics would successfully neutralise the threat of mirror organisms by virtue of their non-stereospecific activity.

There is early evidence that antibiotics would not be effective against mirror bacteria. In the absence of mirror bacteria against which to directly test compounds, two parallel strategies have emerged to determine the likely efficacy of current countermeasures. The first is to synthesise mirror-chirality antibiotics and test them against native bacteria. This strategy relies on the assumption of equivalency between the interaction of mirror antibiotics with native-chirality bacteria and native-chirality antibiotics with mirror bacteria.^10^ While this provides useful experimental evidence, it leverages costly enantiomers of antibiotics and requires “wet lab” resources which constrain the throughput of experiments.

The other strategy is to investigate antibiotic efficacy against mirror bacteria utilising computational chemistry. Thus far, docking and molecular dynamics have been used to investigate the efficacy of amoxicillin against native-chirality and mirror bacterial protein targets from *Chlamydia trachomatis* and *Staphylococcus aureus*.^11,12^ Computational approaches enable investigations into this sensitive topic in a risk-free manner, i.e. without generating any mirror components which may contribute to development of mirror life. In addition, this strategy provides a reproducible and iterative workflow which can be cheaply and rapidly adapted to study a range of targets with minimal resource requirements. This provides a risk-free environment for high-throughput determination of binding efficacy, which can be extrapolated to infer whether antimicrobial activity might be conserved.

In this work, we have leveraged the same computational chemistry approach, extended to a wider range of antibiotics and bacterial protein targets–as outlined in Figure 1. The aims of these experiments were (1) to validate results from other groups, and (2) to determine whether the conclusions were generalisable across antibiotics with differing physicochemical characteristics.

**Figure 1.**
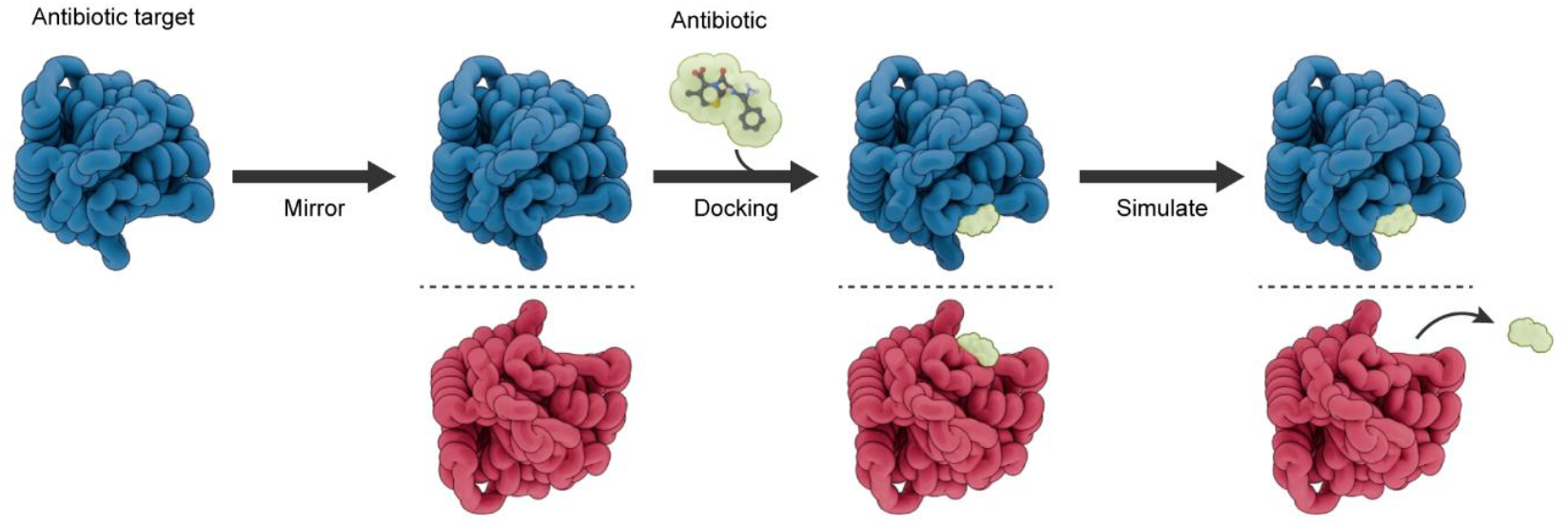
Computational chemistry enables detailed investigations of interactions of antibiotics and their targets, native and mirrored. ***(a)*** Rendered schematic depicting simulation workflow, a selection of antibiotic target proteins from pathogenic species were computationally mirrored, to emulate proteins assembled from D-amino acids in mirror life forms. The native (navy) and mirror (scarlet) proteins were simulated at equilibrium, before performing molecular docking with their constituent antibiotics. Further unrestrained molecular dynamics were performed to determine if the antibiotic would remain in its binding site, suggesting it could inactivate the target protein, and function as an antimicrobial.

**Figure 2.**
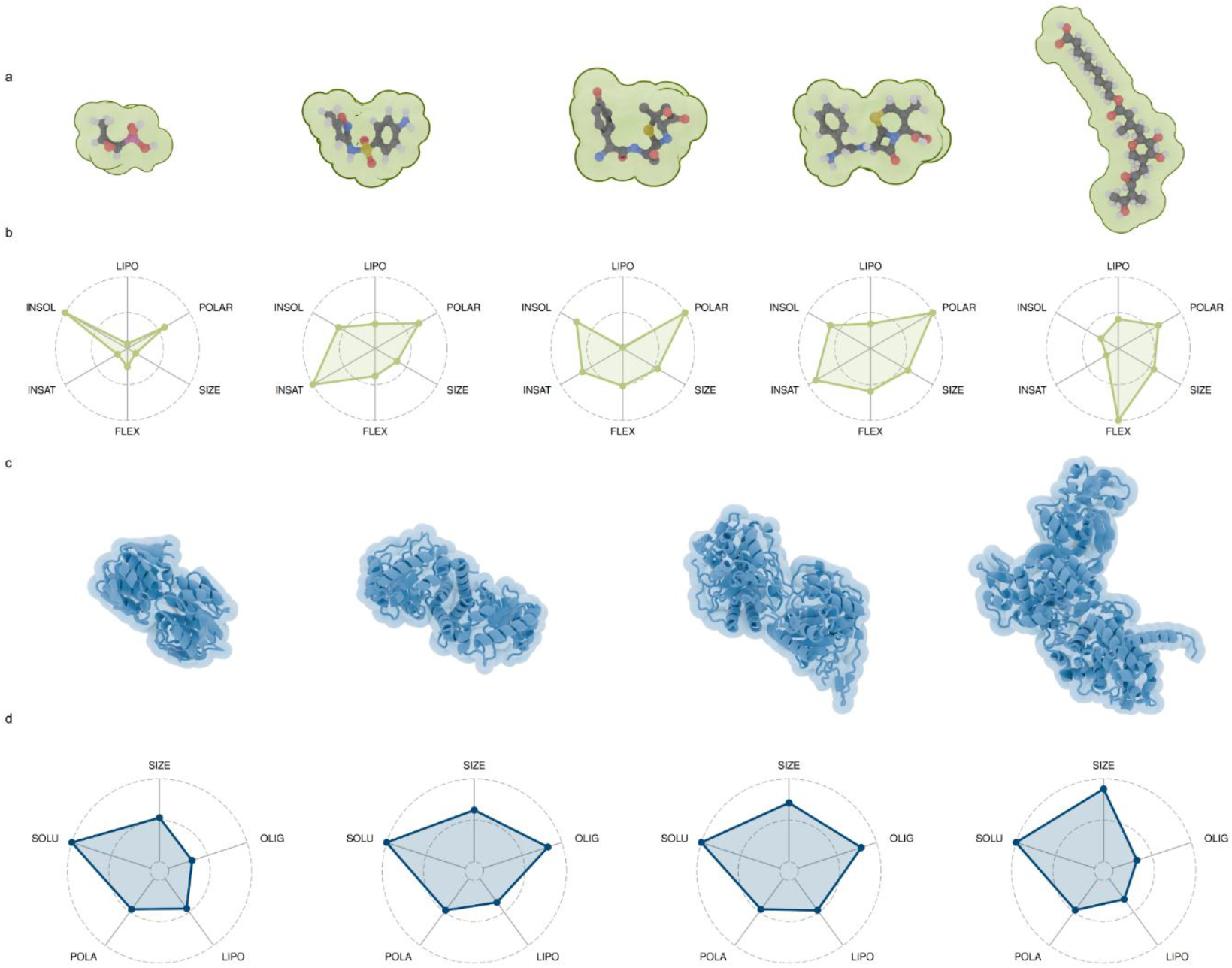
Antibiotics and targets of differing physiochemical and structural properties that were investigated in this work. **(a)** 3D molecular conformers (carbon atoms are depicted grey, hydrogen white, oxygen red, sulfur yellow and nitrogen blue, electrostatic shells are indicated in green) and **(b)** physicochemical properties of antibiotics used in this study, as assessed by SwissADME.^62^ Parameters are lipophilicity (XLogP),^63^ polarity (TPSA),^64^ size (molecular weight), insolubility (ESOL),^65^ insaturation (fraction C_sp3_) and flexibility (number of rotatable bonds). Left to right shows fosfomycin (FFQ, CID:446987), sulfamethoxazole (SMX, CID:5329), amoxicillin (AMX, CID:33613), cephalexin (CPX, CID:27447), and mupirocin (MRC, CID:446596). **(c)** Space-filling and cartoon structures for the 4 proteins investigated in this study. Left to right is UDP-N-Acetylglucosamine 1-Carboxyvinyltransferase from *E. cloacae* (MurA, PDB:3lth)^14^, Dihydropteroate Synthetase from *S. aureus* (DHPS, PDB:1ad1)^15^, the transpeptidase domain of Penicillin Binding Protein 3 from *S. aureus* (PBP3, PDB:6i1f)^16^, and Isoleucyl-tRNA Synthetase from *T. thermophilus* (ileRS, PDB:1jzs)^17^.

## Methods

Protein targets of antibiotics and their mirror images were first simulated at equilibrium. All-atom protein files for a selection of antibiotic targets were downloaded from the protein data bank.^13^ The proteins investigated were UDP-N-Acetylglucosamine 1-Carboxyvinyltransferase from *E. cloacae* (MurA, PDB:3lth)^14^, Dihydropteroate Synthetase from *S. aureus* (DHPS, PDB:1ad1)^15^, the transpeptidase domain of Penicillin Binding Protein 3 from *S. aureus* (PBP3, PDB:6i1f)^16^, and Isoleucyl-tRNA Synthetase from *T. thermophilus* (ileRS, PDB:1jzs)^17^. Mirror image protein structures were generated using a python protocol developed by Pedroni *et al*.^11,12^ Structures were then solvated in SPC/E water and neutralized with Na^+^ and Cl^-^ atoms using *GROMACS* tools (version 2024.3).^18^ Structures were minimised to avoid steric clashes using a steepest descent algorithm for a maximum of 10,000 steps. Systems were then equilibrated to 300 K for 100 ps, using a velocity rescaling thermostat with a 2 ps coupling time. This was followed by isobaric equilibration to 1 bar using isotropic Parrinello-Rahman pressure coupling for 100 ps, also with a 2 ps coupling time. Production simulations were performed for at least 100 ns with two repeats. The CHARMM36 forcefield^19^ was used and electrostatic forces were calculated using the particle-mesh Ewald (PME) algorithm, and a smooth switching algorithm with a 1.2 nm cutoff was used for van der Waals and electrostatic interactions.

Proteins were equilibrated in solution and then docked with their respective antibiotics. Equilibration of native and mirror proteins was assessed by stabilisation of the c-alpha root mean squared deviation (RMSD) and radius of gyration (RoG). Once equilibrated, proteins were docked with antibiotic ligands. Three-dimensional molecular conformations for the ligands were downloaded from PubChem.^20^ Ligands used were sulfamethoxazole (SMX, CID:5329), fosfomycin (FFQ, CID:446987), amoxicillin (AMX, CID:33613), cephalexin (CPX, CID:27447), and mupirocin (MRC, CID:446596). Antibiotic binding sites were identified using the bound antibiotics from the crystallographic data. In the case of DHPS, where there was no bound ligand, a homolog with the correctly bound ligand was used (PDB:3tzf)^21^ and structures aligned using TM-align.^22^ Proteins and their respective ligands were prepared for docking using *Meeko* and docked into a 1.5 nm cube around the binding pocket using *AutoDock Vina*.^23^

Docking poses with the lowest energy minima were taken forward for ligand-bound simulations. Ligand parameters were generated using *cgen-ff*,^24^ and system topologies generated using *gromologist*,^25^ before being prepared, equilibrated, and simulated as described above. Ligand bound trajectories were simulated for at least 100 ns with a minimum of 2 repeats.

Analysis was performed using *GROMACS* tools; *PLIP* was used to identify key binding residues;^26^ *LigPlot+* was used to generate two-dimensional residue interaction maps;^27^ and renders were produced using *VMD*^28^ and Blender. Data analysis and plotting was performed using *R* and *ggplot2*,^29,30^ including the packages *peptides* and *ggradar*.^31,32^

## Results & Discussion

### Enolpyruvyl Transferase (MurA)

Enolpyruvyl transferase (MurA) is the smallest enzyme tested, it acts to catalyse the first step in bacterial cell wall biosynthesis.^33^ MurA is a UDP-N-acetylglucosamine enolpyruvyl transferase, forming UDP-GlcNAc-enolpyruvate and inorganic phosphate from phosphoenolpyruvate (PEP) and UDP-N-acetylglucosamine.^34^ The MurA investigated in this work is a 45.58 kDa monomeric protein of 2 globular domains from the Gram-negative bacteria *Enterobacter cloacae*. This species is one of the most common *Enterobacteriaceae*, and acts as an opportunistic pathogen.^35^ *Enterobacteriaceae* infections can result in bacteremia, sepsis, meningitis, endocarditis and osteomyelitis as well as infections in the urinary and respiratory tracts.^36–38^

When simulated in solution, both the native and mirror forms of MurA rapidly equilibrated. Equilibration status was determined by assessing the RMSD and RoG values of alpha carbons in the protein. Both the native and mirror forms appeared geometrically stabilised after the 100 ns equilibration, consistent with previous reports of simulating proteins this size from crystallography data.^39,40^ After equilibration, native MurA reported an RMSD value of 0.17 ± 0.001, while the mirror form reported an increased 0.28 ± 0.007 nm. Equilibration of the structure was further confirmed by the RoG analysis, which rapidly stabilised to 2.22 ± 0.001 and 2.27 ± 0.03 nm (SI Fig. 1).

MurA is inhibited by the action of the antibiotic fosfomycin. Fosfomycin is a 1.38 kDa broad-spectrum antibiotic. The antibiotic acts as a competitive inhibitor, entering the PEP binding site and alkylating cysteine 115.^41^ Cysteine 115 is located within a highly mobile loop within the enzyme’s active site,^42^ and is an essential component in the catalytic action of MurA.^43^ Alkylation of cysteine 115 also forces the enzyme into an inactive, closed formation, irreversibly blocking peptidoglycan synthesis, which is essential for bacterial survival.^44^

Docking of fosfomycin into the active site of MurA was remarkably similar in the native and mirror forms. In both cases the antibiotic fit into a similar groove in the active site, likely due to the flexibility of the Cys115-containing loops, and demonstrated by the comparable binding energies of -4.13 ± 0.14 and -4.12 ± 0.16 kcal/mol (Fig. 3) for the native and mirror form respectively.

**Figure 3.**
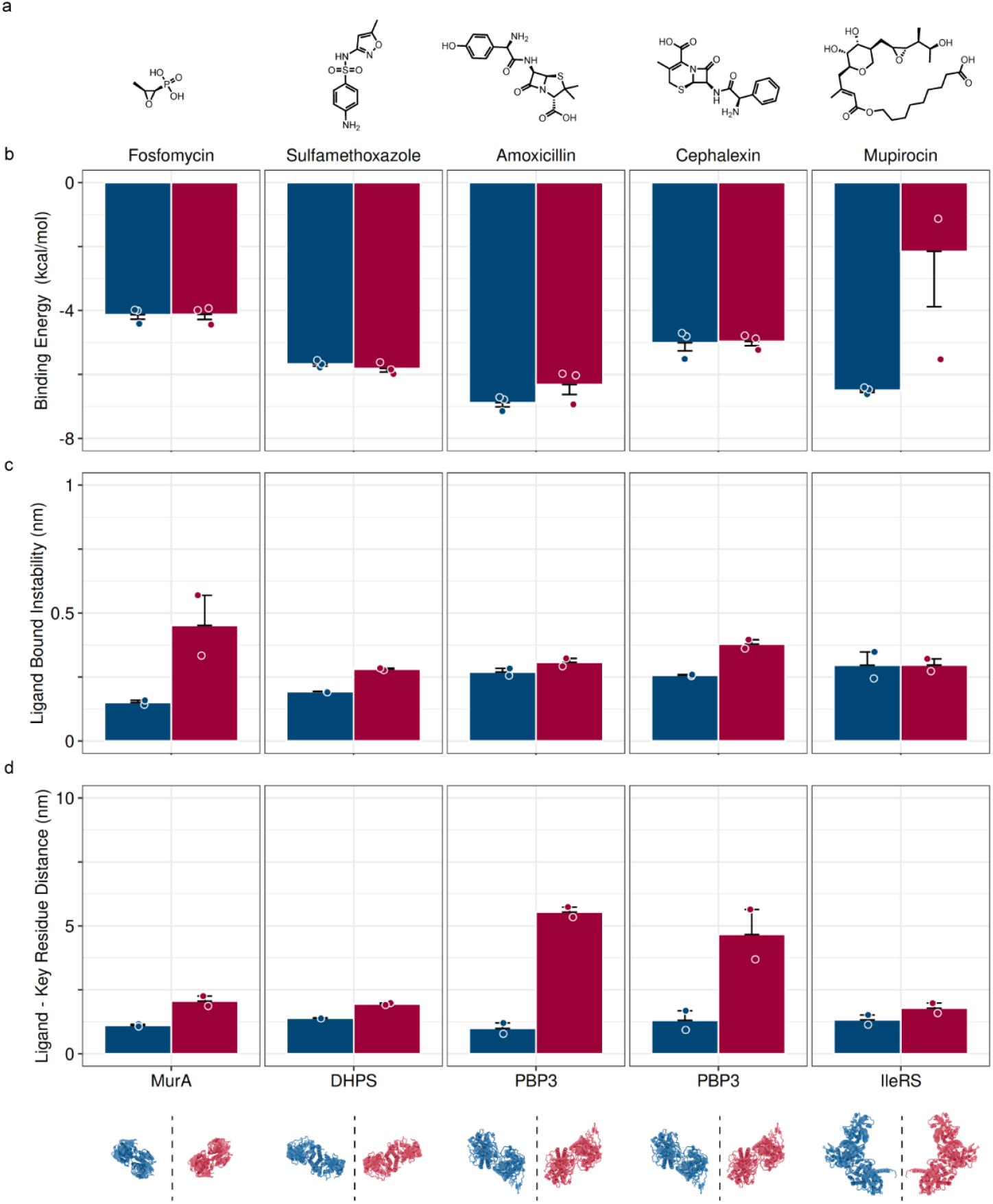
Antibiotics are not effective against mirror targets when compared to their native states, even when well fitted inside target the binding pocket. **(a)** Chemical structures for the antibiotics investigated. **(b)** Binding energies derived when docking antibiotics into the equilibrated native (navy) and mirror (scarlet) targets. **(c)** Protein stability, as measured by alpha carbon RMSD, when simulating post-docking with the bound constituent antibiotic. **(d)** Distance between the antibiotic ligand and the key-residue involved in antibiotic inactivation. Each graph shows the average of at least two repeats, the error bars show the standard error of the mean and the points show the raw data. Key residues were determined using PLIP and LigPlus, as well as literature data and are; MurA-fosfomycin, Cysteine 115; DHPS-sulfamethoxazole, Threonine 51; PBP3-amoxicillin, Serine-320; PBP3-cephalexin, Serine 320; IleRS-mupirocin, Leucine 583.

However, once the antibiotic-bound forms of MurA were freely simulated, the mirror form could stay bound to the antibiotic. While fosfomycin was retained close to the binding site of the mirror form MurA (SI Vid. 2), in both simulations the antibiotic rapidly moved away from the target Cys115 residue (Fig. 4), with an average distance of 2.06 nm ± 0.19, compared to 1.11 ± 0.04 nm in the native form (Fig. 3). In nature, fosfomycin binding is dependent on the initial binding of UDP-Glc-NAC to MurA, causing a conformational shift to enable PEP (or its antibiotic analog fosfomycin) to bind.^45^ In this investigation, this did not appear to be necessary given the tight binding of the native protein to the antibiotic that we observed. Unlike the tight binding of the native form, the distance between the antibiotic fosfomycin and its target Cys115 was large enough that the covalent-bond-mediated inhibition by fosfomycin should not be able to occur, suggesting that the antibiotic may not be effective against mirror form MurA. However, in some simulations the action of fosfomycin caused a significant destabilisation of MurA (Fig. 5, SI Vid. 2), which may also inhibit the enzyme’s function, but stands in stark contrast to the native action of the antibiotic, inducing a closed and non-functional state of the enzyme.

**Figure 4.**
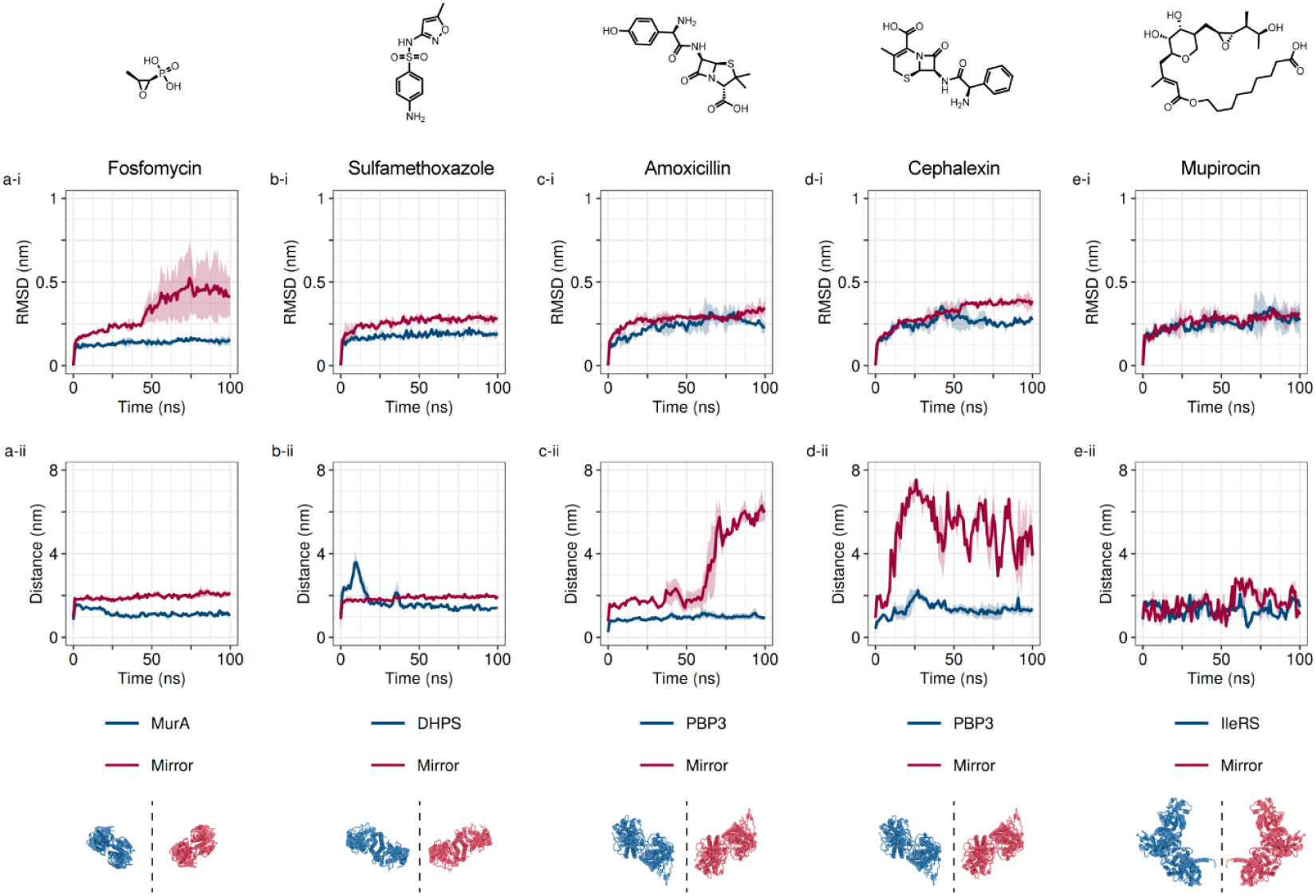
Stability and key residue interactions of antibiotics with their targets and mirrors are antibiotic and target dependent. **(a-i)** Time-averaged alpha carbon RMSD for MurA (navy) and the mirror form (scarlet) once docked with fosfomycin. **(a-ii)** distance between fosfomycin and its key target residue. **(b-i, b-ii)** As above, for DHPS and sulfamethoxazole. **(c-i, cii)** PBP33 and amoxicillin. **(d-i, d-ii)** PBP3 and cephalexin. **(e-i, e-ii)** IleRS and mupirocin. Each trace is the average of at least two repeats, with the transparent ribbon showing the standard deviation.

**Figure 5.**
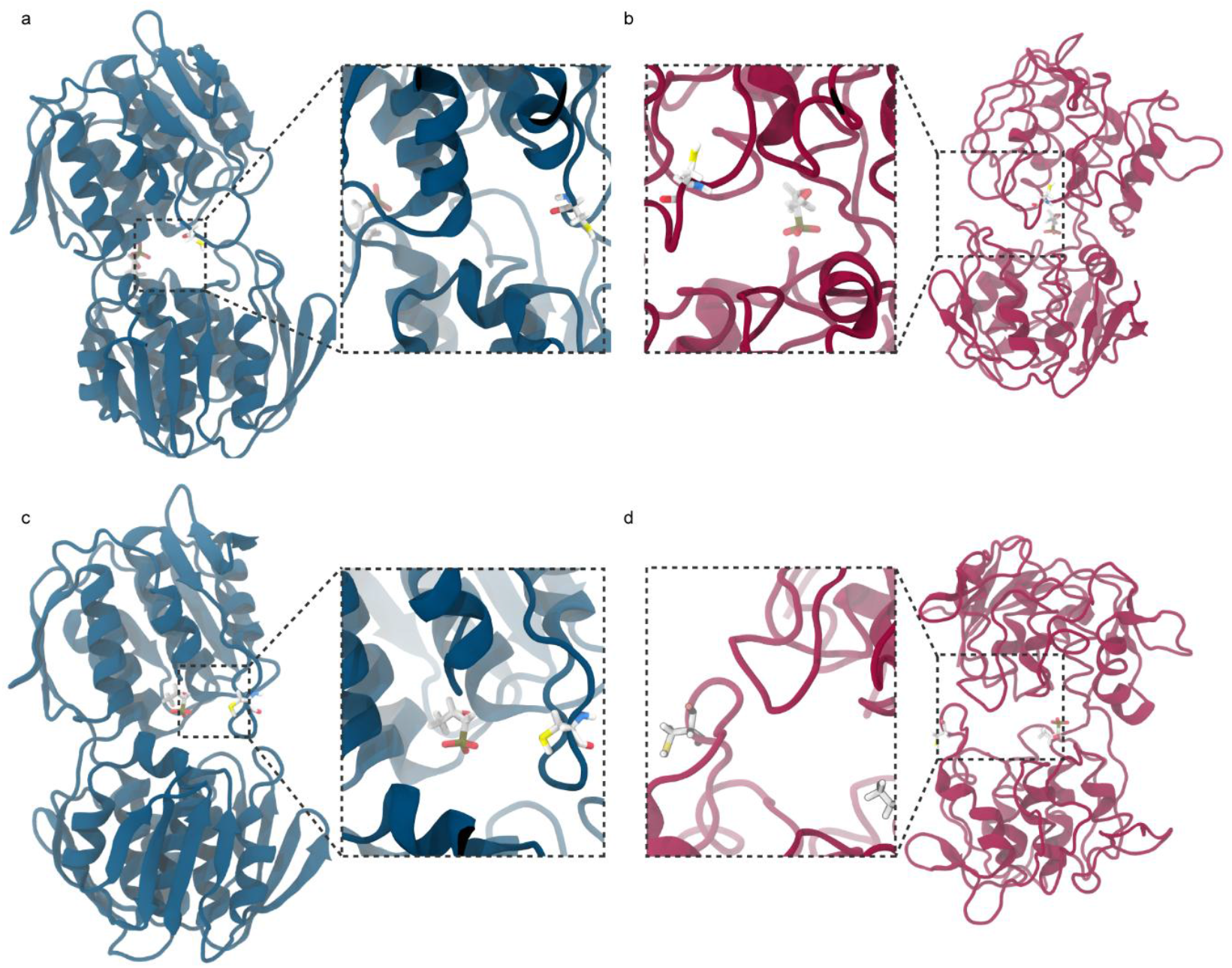
Mirror-MurA retains fosfomycin, but cannot maintain a stable structure. **(a)** Equilibrated native MurA docked with fosfomycin, with cysteine 115 highlighted. **(b)** Equilibrated mirror MurA docked with with fosfomycin. **(c)** Native MurA after 100 ns of unrestrained dynamics with fosfomycin. **(d)** As above, for the mirror form. Proteins are shown in the cartoon representation, native proteins are navy and mirror forms are scarlet. Antibiotics and key residues are depicted as sticks with carbon atoms shown in white, oxygen in red, nitrogen in blue, sulphur in yellow and phosphorus in brown

### Dihydropteroate Synthetase (DHPS)

Dihydropteroate synthase (DHPS) is an enzyme involved in folate biosynthesis, essential in DNA synthesis and repair.^46^ Folate synthesis is a pathway absent in humans, making DHPS a tractable antibacterial target exploited by a range of existing antibiotics. Here we investigated DHPS from *Staphylococcus aureus*,^47^ a human pathogen involved in a range of diseases including pneumonia, sepsis and osteomyelitis, with a well-characterised and extensive antibiotic resistance profile. DHPS is a dimeric protein with two active sites, with flexible loop regions that contain antibiotic target residues threonine 51, phenylalanine 17 and serine 18, which act to catalyse the condensation of 6-hydroxymethyl-7,8-dihydropterin-pyrophosphate and *p*-aminobenzoic acid to produce folate.^48^

Equilibration simulations demonstrated rapid plateauing of both native and mirror form DHPS, suggesting geometric stability (SI Fig. 1). Average RMSD values for DHPS in the absence of antibiotics were 0.21 ± 0.006 nm for the native form and 0.29 ± 0.001 nm for the mirror form. RoG was also used to confirm equilibration, with both forms demonstrating stability with average values of 2.68 ± 0.008 and 2.65 ± 0.006 (SI Fig. 1). The RMSD values we obtained are slightly higher than previously reported.^49^ This is likely due to the use of different forcefields and omission of stabilising magnesium ions.

DHPS is inhibited by the antibiotic sulfamethoxazole. Sulfamethoxazole is a competitive inhibitor which prevents *p*-aminobenzoic acid from accessing the binding site.^50^ This inhibition is reversible, acting to sterically inhibit the natural substrate as opposed to inactivating key residues. However mutation of key residues such as T51M can inhibit sulfamethoxazole binding, while still retaining enzyme function, resulting in clinical resistance.^50^

Docking of sulfamethoxazole into the native and mirror forms of DHPS showed very consistent results (Fig. 4). Average binding energies were -5.67 ± 0.07 and -5.81 ± 0.11 kcal/mol (Fig. 3) for the native and mirror form respectively. In both cases the antibiotic was able to easily enter the active site of the protein.

Once bound to sulfamethoxazole, both native and mirror forms showed increased stability over the unbound forms (Fig. 3). Average RMSD values for the native and mirror forms were 0.19 ± 0.002 and 0.28 ± 0.004 nm respectively, suggesting stabilisation of both forms, although the native state was marginally more stable. In both cases the antibiotic is well retained in the active site, while it is not necessarily always close to the key residue selected (Thr51), this is not necessary for effective inhibition. Average distances between Thr51 and sulfamethoxazole were 1.4 ± 0.02 nm and 1.95 ± 0.04 nm for the native and mirror form respectively. In the native state, some re-organisation of the antibiotic is visible (SI Vid. 3) which is not visible in the mirror form, allowing the antibiotic closer to Thr51. However it is likely that in both cases the activity of the enzyme would be sterically impeded.

### Penicillin-Binding Protein 3 (PBP3)

Penicillin-Binding Protein 3 (PBP3) is an essential enzyme involved in peptidoglycan crosslinking during cell division.^51^ The PBP3 form tested was from *Chlamydia trachomatis*, an obligate intracellular pathogen commonly involved in urogenital infections, which can result in chronic inflammation and scarring. PBP3 is a 73.67 kDa enzyme, however only the 36.5 kDa TP domain was simulated.^16^ The TP domain was selected both to reduce compute requirements, and to confirm the simulation methodology used here gave results consistent with that of previous work from Pedroni *et al*.

As with the previous proteins, the small TP domain rapidly reached geometric stability (SI Fig. 1). Average RMSD values for PBP3 in the absence of antibiotics were 0.25 ± 0.02 for the native form and 0.36 ± 0.01 for the mirror form, consistent with the work from Pedroni *et al*. Further demonstration of equilibration came from the stable RoG of the native form (1.98 ± 0.001), which was consistent with that of the mirror form (1.99 ± 0.01), which had fully stabilised after the 100 ns equilibration, despite briefly spiking at 50 ns (SI Fig. 1).

PBP3 was assessed with 2 different antibiotics, the widely used β-lactams amoxicillin and cephalexin. Both antibiotics inhibit cell maturation and differentiation.^52^ The antibiotics form a covalent adduct with serine 320 in in the PBP active site, inhibiting peptidoglycan cross-linking.^53^

Docking PBP3 in complex with amoxicillin and cephalexin showed comparable binding strengths. In all cases the antibiotic could fit into the active site of both the native and mirror form. The average binding energies of the top 3 docking poses were -6.88 ± 0.13 and -6.31 ± 0.31 kcal/mol for the native and mirror PBP3 binding to amoxicillin, and -5.01 ± 0.26 and -4.96 ± 0.14 kcal/mol for cephalexin (Fig. 3).

When simulated with the bound antibiotics, both the native and mirror proteins were stable, but the mirror form was unable to retain the antibiotics. Protein stability was consistent across repeat simulations with RMSD values of 0.27 ± 0.01 and 0.31 ± 0.02 nm for the native and mirror forms complexed to amoxicillin (respectively) and 0.26 ± 0.003 and 0.38 ± 0.02 nm for the native and mirror forms complexed to cephalexin (respectively) (Fig. 3). Despite this stability, the distance between the antibiotics and their target residue Ser320 demonstrated the ineffectiveness of the antibiotics against the mirror form bacterial target. In the native forms amoxicillin was retained with 0.99 ± 0.2 nm of Ser320, while cephalexin was 1.31 ± 0.38 nm (Fig. 4). However, the mirror forms saw the antibiotics exit the binding site within 10 ns for cephalexin and 60 ns for amoxicillin, with final distances being 5.54 ± 0.2 nm for amoxicillin and 4.67 ± 1 nm for cephalexin (Fig. 4). This strongly suggests that the antibiotics would be ineffective – although it may be feasible that amoxicillin could have some long-lasting inhibitory action in the 50 ns it was retained in the binding site, this is a very short timeframe in which to act and it is evidently not as well retained in the native form (Fig. 6).

**Figure 6.**
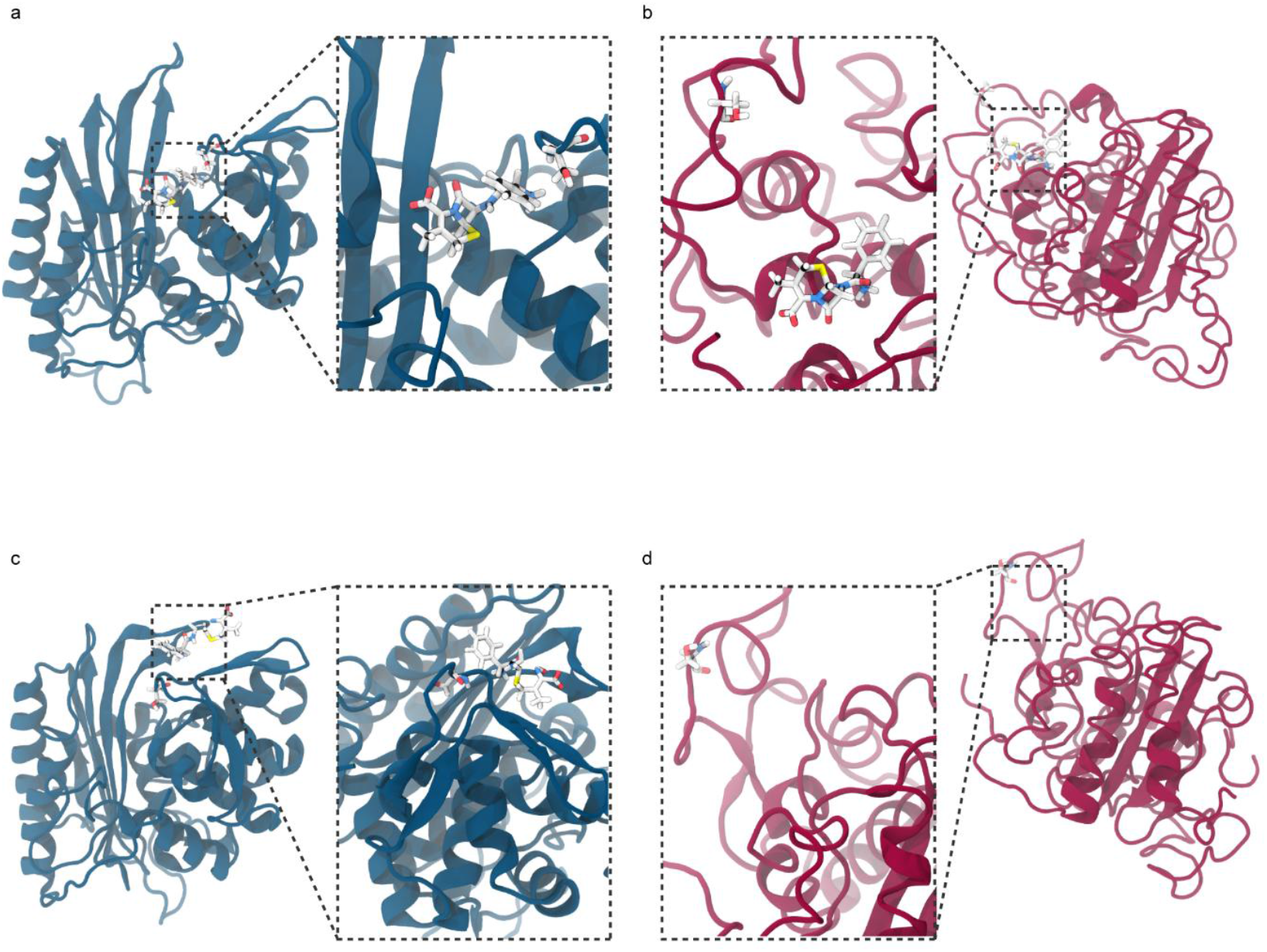
Mirror PBP3 cannot retain cephalexin within the binding pocket. **(a)** Equilibrated native PBP3 docked with cephalexin, with serine 320 highlighted. **(b)** Equilibrated mirror PBP3 docked with cephalexin. **(c)** Native PBP3 after 100 ns of unrestrained dynamics with cephalexin. **(d)** As above, for the mirror form. Proteins are shown in the cartoon representation, native proteins are navy and mirror forms are scarlet. Antibiotics and key residues are depicted as sticks with carbon atoms shown in white, oxygen in red, nitrogen in blue, sulphur in yellow.

**Figure 7.**
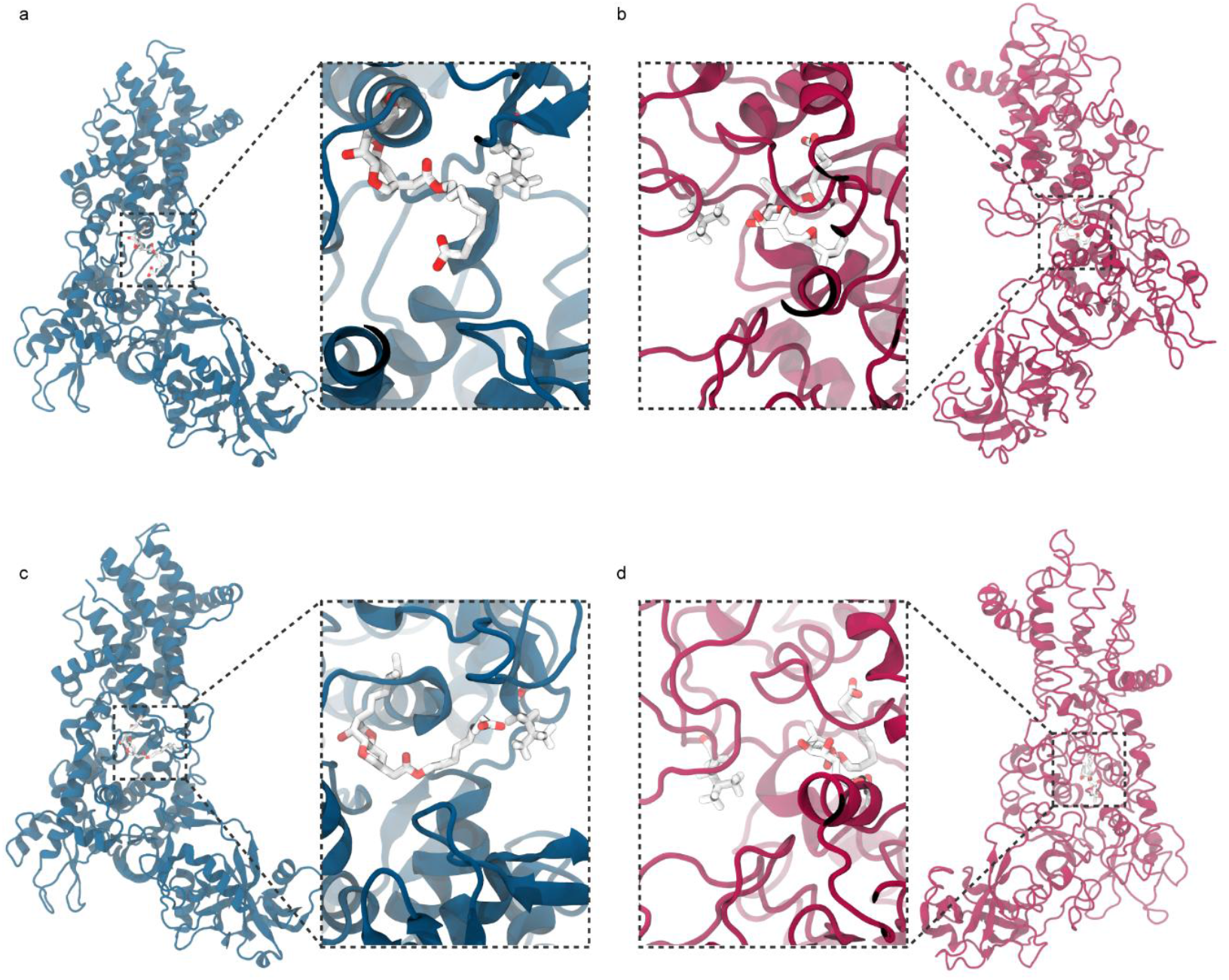
Flexible thermophilic pathogen targets were better able to retain their antibiotic ligand. **(a)** Equilibrated native IleRS docked with mupirocin, with leucine 583 highlighted. **(b)** Equilibrated mirror IleRS docked with mupirocin. **(c)** Native IleRS after 100 ns of unrestrained dynamics with mupirocin. **(d)** As above, for the mirror form. Proteins are shown in the cartoon representation, native proteins are navy and mirror forms are scarlet. Antibiotics and key residues are depicted as sticks with carbon atoms shown in white, oxygen in red, nitrogen in blue, sulphur in yellow.

The computational process used to form the mirror proteins could also be used on the antibiotics, so the mirrored form of amoxicillin was assessed against native and mirror PBP3. When tested our results were comparable with previous work,^12^ where the mirrored antibiotic could be retained in the mirrored protein, although it was not a tight binding and a decrease in stability was detected (SI Fig. 1). Building on previous work we also investigated the efficacy of the mirror antibiotic on native PBP3, in which predictably the mirror antibiotic could not be retained in the active site of the native protein (SI Vid. 5-8).

### Isoleucyl-tRNA Synthetase (IleRS)

isoleucyl-tRNA synthetase (IleRS) is an essential aminoacyl tRNA synthase enzyme. It acts to assemble isoleucine-tRNA complexes involved in protein synthesis by transferring aminoacyl adenylate to the 3’ terminal adenosine of tRNA.^54^ The IleRS investigated here is from the thermophilic eubacterium, *Thermus thermophilus*, selected for the high quality crystallography data in complex with the antibiotic mupirocin.^17^ While *T. thermophilus* is not a human pathogen, tRNA synthesis is highly conserved and there is no significant difference in the amino acid sequence between catalytic domains of IleRS proteins from different kingdoms.^54^

As the largest protein studied, IleRS showed some of the highest RMSD and RoG values, but still appeared to rapidly reach geometric equilibrium. The 95.31 kDa protein appeared to reach stability in both native and mirror forms within the first 10 ns and RMSD and RoG were consistent from that time onwards. Average RMSD for the native IleRS was 0.29 ± 0.02 nm, and the mirror form again showed slightly higher RMSD than the native state at 0.36 ± 0.001 nm (SI Fig. 1). Despite the small change in stability induced by computational mirroring, the RoG values were consistent with both native and mirror forms showing 3.46 ± 0.002 (SI Fig. 1).

IleRS is inhibited by the action of the antibiotic mupirocin. Mupirocin is an analog of the isoleucyl-adenylate substrate, competitively excluding it from the active site and disabling protein synthesis.^54^ Mupirocin is believed to bind to leucine 583 in the active site, as its substitution in eukaryotic IleRS confers resistance to mupirocin.^54,55^

The docking of mupirocin into the active site of IleRS showed the largest difference between the native and mirror state of any of the proteins tested. Native IleRS showed highly consistent docking, with a binding energy of -6.5 ± 0.06 kcal/mol. However the mirrored form was highly inconsistent, the top docking pose had a comparable binding energy to the native (Fig. 3), but the rest were bound significantly less strongly, resulting in an average binding energy of -2.2 ± 1.7 kcal/mol. Despite the binding energy being significantly reduced compared to the native protein, it was still spontaneous, suggesting binding may occur.

IleRS from *T. thermophilus* is the only protein tested to show comparable binding and stability to its cognate antibiotic between the native and mirrored form. The antibiotic bound trajectories had comparable RMSDs (Fig. 3) of 0.3 ± 0.05 and 0.3 ± 0.02 nm for the native and mirror states respectively, nearly identical to the unbound RMSD. The average distance between the mupirocin and its target residue, Leu583, was 1.33 ± 0.19 and 1.79 ± 0.2 nm in the native and mirror state respectively (Fig. 3), with the antibiotic staying tightly bound into the active site in both the mirror and native forms (Fig. 4, Fig. 7). This may be due to the highly flexible nature of both mupirocin and IleRS. Mupirocin has a long hydrocarbon tail, conferring flexibility on the molecule; complementarily, IleRS is highly flexible, a known feature of proteins from thermophilic species (Fig. 2).^56^ However, the variable docking scores suggest that it may be unlikely for the antibiotic to spontaneously enter the binding site, even if it can be retained once there. It is interesting that the only instance of possible antibiotic effectiveness against both native and mirror forms in our work comes from a species that is not pathogenic to humans. We believe that this derives from the extremophilic nature of the protein, unlike which

## Conclusions

In most cases, antibiotics were not able to remain in close proximity to binding pockets of mirror forms of bacterial proteins – unlike native forms. From this, it may be inferred that the antibiotics tested would fail to exert an antimicrobial effect on potential future mirror bacteria. While most antibiotics were shown to be ineffective, antibiotics which act through steric hindrance (as opposed to covalent modification) may have more potential against mirror life, as the conditions for effectiveness are often less spatially stringent. This is likely to depend on the size and flexibility of the binding pocket. This is worrying as antibiotics which work by covalent modification are often more potent and selective antibiotics,^57^ as well as one of the few types of antibiotics in which we continue to see discoveries.^58^ Some of the most historically important and widely used antibiotics, such as the β-lactams, work through covalent modification of target proteins.^59^ Antibiotics which are highly flexible, or target flexible proteins, are likely to be more effective against mirror life. This is a positive effect that warrants further investigation as part of any future anti-mirror drug development efforts. These could draw on existing drug development strategies, as many antibiotics such as linezolid and related oxazolidinones have been optimised by increasing their rotational freedom.^60^ Further, many more modern antibiotics (especially peptide mimics)^61^ are highly flexible and may as a result be more effective against mirror life. Regardless, the data presented here suggests that, considering the current crises of effective antibiotics, adding mirror life bacteria into the fray represents a serious biosecurity threat which warrants significant further investigation.

## Contributors and sources

Paul-Enguerrand Fady: Conceptualization, Project administration, Writing - Original Draft, Writing - Review & Editing, Funding acquisition

Jonah L Ciccone: Investigation, Data generation, analysis & curation, Methodology, Visualization, Writing - Original Draft, Writing - Review & Editing,

## Acknowledgements

The authors acknowledge the use of the UCL Kathleen High Performance Computing Facility (Kathleen@UCL) and associated support services in the completion of this work. The authors also acknowledge Ana Leonescu for insightful input regarding the chemistry of small molecule antibiotics.

## Data availability

All trajectories are available upon request.

---

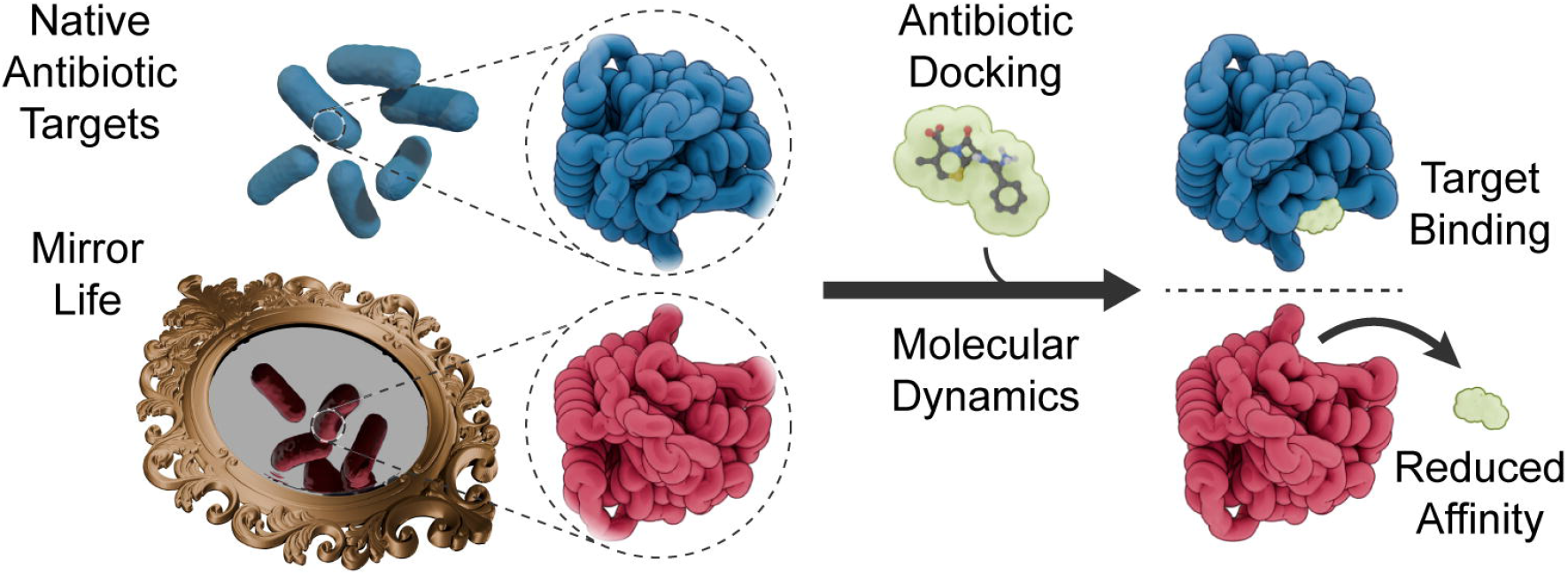

