## Supplementary material for "Molecular dynamics simulations demonstrate reduced antibiotic affinity to mirror bacterial targets": SI

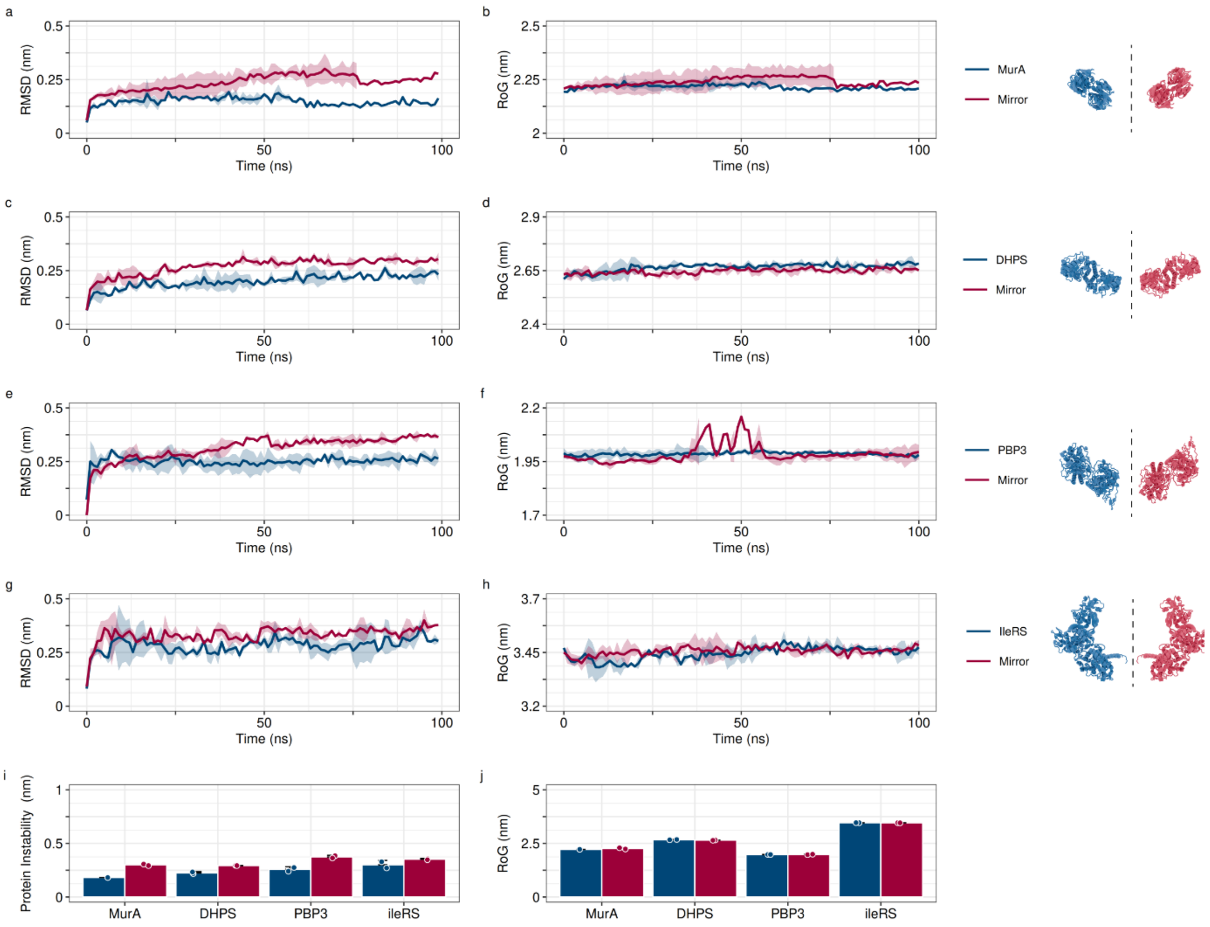


SI Figure 1. **Native and mirror form proteins reached geometric equilibrium after 100 ns. (a)** Time-averaged alpha carbon RMSD and **(b)** RoG for MurA, **(c-d)** DHPS, **(e-f)** PBP3 and (**g-h)** IleRS. Native proteins are shown in navy and mirror in scarlet. Each trace is an average of 2 repeats and the coloured ribbon shows the standard deviation. **(i)** Average RMSD and **(j)** RoG from the final 30 ns of each trajectory. Each graph shows the average of at least two repeats, the error bars show the standard error of the mean and the points show the raw data.

| Protein | Ligand | Enantiomer | RMSD(nm) | S.E.M. |
|---|---|---|---|---|
| MurA | Unbound | Native | 0.17 | 0.001 |
| MurA | Unbound | Mirror | 0.28 | 0.007 |
| DHPS | Unbound | Native | 0.21 | 0.006 |
| DHPS | Unbound | Mirror | 0.29 | 0.001 |
| PBP3 | Unbound | Native | 0.25 | 0.021 |
| PBP3 | Unbound | Mirror | 0.36 | 0.013 |
| IleRS | Unbound | Native | 0.29 | 0.02 |
| IleRS | Unbound | Mirror | 0.35 | 0.001 |
| MurA | Fosfomycin | Native | 0.15 | 0.009 |
| MurA | Fosfomycin | Mirror | 0.45 | 0.118 |
| DHPS | Sulfamethoxazole | Native | 0.19 | 0.002 |
| DHPS | Sulfamethoxazole | Mirror | 0.28 | 0.004 |
| PBP3 | Amoxicillin | Native | 0.27 | 0.014 |
| PBP3 | Amoxicillin | Mirror | 0.31 | 0.015 |
| PBP3 | Cephalexin | Native | 0.26 | 0.003 |
| PBP3 | Cephalexin | Mirror | 0.38 | 0.017 |
| IleRS | Mupirocin | Native | 0.3 | 0.052 |
| IleRS | Mupirocin | Mirror | 0.3 | 0.024 |

SI Table 1. Time-averaged Root Mean Squared Deviation (RMSD) in nm for carbon alpha atoms of each antibiotic target protein taken from equilibrated regions of the trajectory. Each value is the average of two independent simulations, and standard error of the mean is given. Data is given for simulations performed in the presence and absence of constituent antibiotics.

| Protein | Enantiomer | RoG(nm) | S.E.M. |
|---|---|---|---|
| MurA | Native | 2.22 | 0.001 |
| MurA | Mirror | 2.27 | 0.03 |
| DHPS | Native | 2.68 | 0.008 |
| DHPS | Mirror | 2.65 | 0.006 |
| PBP3 | Native | 1.98 | 0.001 |
| PBP3 | Mirror | 1.99 | 0.011 |
| IleRS | Native | 3.46 | 0.002 |
| IleRS | Mirror | 3.46 | 0.002 |

SI Table 2. Time-averaged Radius of Gyration (RoG) in nm for carbon alpha atoms of each antibiotic target protein taken from equilibrated regions of the trajectories. Each value is the average of two independent simulations, and standard error of the mean is given. Data is taken from simulations performed in the absence of antibiotics.

| Protein | Ligand | Enantiomer | BindingEnergy(kcalmol$^{-1}$) | S.E.M. |
|---|---|---|---|---|
| PBP3 | Amoxicillin | Native | -6.88 | 0.134 |
| PBP3 | Amoxicillin | Mirror | -6.31 | 0.312 |
| PBP3 | Cephalexin | Native | -5.01 | 0.255 |
| PBP3 | Cephalexin | Mirror | -4.96 | 0.136 |
| IleRS | Mupirocin | Native | -6.5 | 0.063 |
| IleRS | Mupirocin | Mirror | -2.15 | 1.731 |
| DHPS | Sulfamethoxazole | Native | -5.67 | 0.067 |
| DHPS | Sulfamethoxazole | Mirror | -5.81 | 0.106 |
| MurA | Fosfomycin | Native | -4.13 | 0.141 |
| MurA | Fosfomycin | Mirror | -4.12 | 0.162 |

SI Table 3. Average binding energies for each antibiotic docked into the binding site of their target protein. Each value is the average of the top three docking poses and the standard deviation is given.

| Protein | Ligand | Enantiomer | KeyResidue | Distance(nm) | S.E.M. |
|---|---|---|---|---|---|
| MurA | Fosfomycin | Mirror | CYS_115 | 1.11 | 0.044 |
| MurA | Fosfomycin | Native | CYS_115 | 2.06 | 0.193 |
| DHPS | Sulfamethoxazole | Mirror | THR_51 | 1.4 | 0.015 |
| DHPS | Sulfamethoxazole | Native | THR_51 | 1.95 | 0.039 |
| PBP3 | Amoxicillin | Mirror | SER_320 | 0.99 | 0.211 |
| PBP3 | Amoxicillin | Native | SER_320 | 5.54 | 0.198 |
| PBP3 | Cephalexin | Mirror | SER_320 | 1.31 | 0.375 |
| PBP3 | Cephalexin | Native | SER_320 | 4.67 | 0.976 |
| IleRS | Mupirocin | Mirror | LEU_583 | 1.33 | 0.189 |
| IleRS | Mupirocin | Native | LEU_583 | 1.79 | 0.194 |

SI Table 4. Time-averaged distance in nm between each antibiotic and a key target residue within the binding site of its target protein, taken from equilibrated regions of the trajectory. Each value is the average of two independent simulations, and standard error of the mean is given.

SI Video 1. Representative trajectory of MurA after docking with fosfomycin (100 ns)

SI Video 2. Representative trajectory of mirror for MurA after docking with fosfomycin (100 ns)

SI Video 3. Representative trajectory of DHPS after docking with sulfamethoxazole (100 ns)

SI Video 4. Representative trajectory of mirror for DHPS after docking with sulfamethoxazole (100 ns)

SI Video 5. Representative trajectory of PBP3 after docking with amoxicillin (100 ns)

SI Video 6. Representative trajectory of mirror for PBP3 after docking with amoxicillin (100 ns)

SI Video 7. Representative trajectory of PBP3 after docking with cephalexin (100 ns)

SI Video 8. Representative trajectory of mirror for PBP3 after docking with cephalexin (100 ns)

SI Video 9. Representative trajectory of IleRS after docking with mupirocin (100 ns)

SI Video 10. Representative trajectory of mirror for IleRS after docking with mupirocin (100 ns)
